# Dataset on Emotion with Naturalistic Stimuli (DENS) on Indian Samples

**DOI:** 10.1101/2021.08.04.455041

**Authors:** Sudhakar Mishra, Mohammad Asif, Narayanan Srinivasan, Uma Shanker Tiwary

**Affiliations:** Indian Institute of Information Technology Allahabad, Information Technology, Prayagraj, 211012, India; Indian Institute of Technology-Kanpur, Cognitive Science, Kanpur, 208016, India; University of Allahabad, Centre of Behavioural and Cognitive Sciences, Prayagraj, 211002, India

## Abstract

Emotions are constructed and emerge through the dynamic interaction of multiple components. It is difficult to capture the dynamics using static or artificial stimuli. Hence, there is a need for an experiment paradigm using ecologically valid film stimuli. The data set described in this work results from an attempt to capture felt emotional experience at a particular point in time using physiological measures like EEG, ECG and EMG as well as self-reported scales. Sixteen emotional film stimuli were used from the film stimuli dataset validated in the Indian population. Participants self-reported the felt emotional category. Both the raw and pre-processed data are provided along with the pre-processing pipeline. The paradigm we have adopted is new which we have termed as Emotional Event Marker Paradigm (EEMP). Hence, the dataset has unique information about temporal markers of emotional experiences while watching the film stimuli, which is not available with any data to date. It is the first EEG data with emotional film stimuli on the Indian population. This data can be utilized to study dynamic activation and connectivity in a whole-brain source localization study, understand the mutual interactions between the central and autonomic nervous system, understand temporal hierarchy using multi-resolution tools, and perform machine learning-based classification and complex networks analysis associated with emotions.

## Background

Emotions are the mental phenomena which affect almost every faculty of the Brain including decision making^1^, perception^2^, problem-solving^3^, learning^4^, memory^4^, and conscious selfhood^5^. Yet, it is largely a mystery that how emotions are processed in the brain and how these processes can be tapped to achieve an efficient functioning brain. Although western society has made some progress in understanding the emotion dynamics in the Western context, the understanding of emotion dynamics in the Indian context is largely untouched.

Western datasets like MAHNOB-HCI used multimodal affective stimuli and recorded emotional responses of subjects^6^. In addition to EEG the dataset had other modalities including face videos, audio signals, eye gaze data, and peripheral nervous system physiological signals. Twenty-seven subjects self-reported their emotional response in terms of valence, arousal, dominance, predictability and emotion words. Though, the primary aim of the author was to avail a dataset for the emotion recognition. Hence, they didn’t had time specific information about the emotional experience. Similar, idea is applied to the creation of DEAP data^7^, DECAF data^8^, AMIGOS data^9^ and ASCERTAIN data^10^. Emotion recognition has importance in its own right but it lacks in creating the understandin regarding the neural ldynamics of emotions. Dataset with multimodal stimuli has its advantage in term of ecological validity but it also carries downside of lack of localization of emotional experience.

In our experiment, we tried to bridge this trade-off by utilizing the advantages of multimodal film stimuli along with addressing the problem of temporal localization of emotional experience. We found no other dataset on emotions with naturalistic and real life resembling dynamic stimuli which also provide the temporal marker where the participants had felt any emotion. Hence, we claim that the dataset which we have recorded and describing in this work has unique, novel, and realistic information about emotional experience.

Our data could add a valuable contribution in the emotion research in the following ways.

- Emotions are constructed and experienced in a particular context. Hence, there is a need to study emotion processing using more naturalistic paradigms to understand the emotional dynamics with its multi-component constituents. We collected data in a more ecologically valid experimental setting using naturalistic emotional stimuli.
- This data could be utilized by researchers trying to understand emotion dynamics whose efforts are hindered by the use of static or less ecologically valid stimuli without context.
- Complex networks analysis and dynamic tracking of causality can be performed on this data. In addition, machine learning-based classification can determine the characteristic statistical signature of emotions and their processing.
- Since the information about the time duration when participants felt any emotion is provided, our data is not contaminated by the mind-wandering activity.

## 1 Method

In this paper we are presenting an EEG dataset along with recording of peripheral physiological signals. In addition, the 44-item Big Five Inventory (BFI;^11^) was used to measure the five broad personality traits. All items were evaluated on a 5-point Likert scale, ranging from “strongly disagree” to “strongly agree.”

### 1.1 Big-5 Personality Traits

The Big-five personality traits that are observed among individuals are Openness, Conscientiousness, Extraversion, Agreeableness, and Neuroticism (OCEAN). The reaction of an individual in any given situation depends on these personality traits. According to psychologists the personality of a human can be deciphered using these five basic personality traits. The factors included in OCEAN helps in identifying the behaviour of a person because the encapsulate the qualities of a person. Reportedly, across cultures these five major personality traits describes the individual’s behaviour. One personality trait dominates in each one of us that reflects our character. The individual personality traits are described as follows:

1. **Neuroticism:**Least emotionally stable are categorized in the neuroticism trait. People with neuroticism trait get instigated by little things. They can easily get upset. They are less resilient to negative mood and have difficulty in managing stress. Due to the low emotional stability a person with neuroticism trait tend to see negative side of the things and is prone to negative mood related problems, for example, anxiety, depression.
2. **Extraversion:**Person with high social skill like friendly nature, interaction with people and engaging in social or group events (e.g. parties) pertain to extraversion trait. They like to be around people, hate being alone and engage in deep discussions. They tend to be talkative, seekers and enthusiastic about any social event.
3. **Agreeableness:**People with agreeable personality trait are polite and compassionate in nature. Generally, people with this trait are good hearted, polite, courteous, trustworthy and extremely cooperative. They are respectful to their colleagues and have have high empathy in nature.
4. **Conscientiousness:**Conscientiousness personality characterise those people who are self-disciplined and dutiful. The characteristic of being well-organized, responsible, stick to plan, and goal orientation adhere to conscientiousness trait. People with conscientiousness trait try to realize the impact of their actions beforehand and like to execute their plans step by step. However, their plans and actions are confined to the boundary of moral decency and ethical believes. Punctuality, reliability, time management and sticking to the rules are their primary concerns. For instance, their homes are clean and organized. Free from any short of clutter. Their day to day necessities are arranged properly rather than haphazard these stuffs in the room.
5. **Openness:**People who display openness personality are receptive of new ideas, open minded, adventurous, open to new experience, open to enhance their knowledge, enjoy creativity, artistic. On the contrary, low openness are characterised with conservative and unacceptable to liberal views.

### 1.2 DSM-V Level-1

Using DSM-5 level 1 scale^12^, the mental health of participants are measured. The measurement consists of 23 questions encompassing the assessment of 13 psychiatric domains including sleep problems, depression, somatic symptoms, repetitive thoughts and behaviors, substance use, anger, suicidal ideation, dissociation, personality functioning, psychosis, mania, memory and anxiety. This scale bothers considers the influence of the above domains in behaviour during the past two weeks.

Left to right ratings are marked from 0 to 5, respectively, none or not at all; slight or rare, less than a day or two; mild or several days; moderate or more than half the days; and severe or nearly every day).

### 1.3 Participants

Although participants were given presentations about the experiment, they were also awarded with grade credit for their participation. Forty-three participants has participated in the study. These participants were all master’s level student from Indian Institute of Information Technology Allahabad. Data of three participants with excessive movement was removed. The participants age were varied from 18 to 30 years with the mean age = 23.3 ±1.4 years and females = 3. Ratings on DSM Axis I were used as inclusion and exclusion criteria. However, no participant were removed based on DSM ratings. All participants provided written informed consent and were compensated for their time. Institutional review board of University of Allahabad reviewed the experiment and granted permission with the protocol (protocol code 2017-100 approved on Dec 8, 2017).

### 1.4 Stimuli

For the EEG experiment 16 emotional stimuli were selected from the affective multimedia stimuli dataset validated on Indian samples^13^. The assigned emotional categories to these 16 stimuli (assigned in the stimuli validation work), includes Adventurous (ID:199), Afraid (ID:214), Alarmed (ID:10), Amused (ID:99), Angry (ID:198), Aroused (ID:54), Calm (ID:40), Disgust (ID:26), Enthusiastic (ID:92), Excited (ID:152), Happy (ID:109), Joyous (ID:201), Melancholic (ID:51), Miserable (ID:210), Sad (ID:113), and Triumphant (ID:67). See the Appendix-1 for the stimuli table. The time duration of each stimulus is 60 seconds. We also validated two non-emotional stimuli with the same 60 seconds duration.

The validation of non-emotional stimuli was done with 15 participants. Each participants were shown the same eight non-emotional videos. Two of the eight stimuli were not rated by any participant in any emotional category. In addition, the valence and arousal values for these non-emotional stimuli were around five as shown in the table-2.

**Table 1.** Multi-modal fatasets in the emotion literature. Most of the datasets are created with the goal of pattern recognition in mind. Abbr: I - Individual; G - Group.

| Database | Part | Individual vs Group | Purpose | Modalities |
| --- | --- | --- | --- | --- |
| MAHNOB-HCI<sup>6</sup> | 27 | I | Implicit Tagging and Recognition of Emotion | Peripheral: ECG, Respiration Amplitude, GSR & Skin Temperature; Central: EEG; Audio-Visual and Eye Gaze |
| DEAP<sup>7</sup> | 32 | I | Peripheral and EEG signals for Implicit affective tagging | Peripheral: Respiration Amplitude, Blood Volume, EMG, EOG, Skin Temperature, GSR; Central: EEG; Visual for 22 participants. |
| DECAF<sup>8</sup> | 30 | I | Affective Annotation | Peripheral: Trapezius EMG, hEOG, ECG; Central: MEG; Facial Expression |
| ASCERTAIN<sup>10</sup> | 58 | I | Probing Affect & Personality | Peripheral: GSR, ECG; Central: EEG; Visual |
| AMIGOS<sup>9</sup> | 40 | I & 4 people Group | Recognition of Mood, Personality, Affect & Social Context Recognition | Peripheral: ECG and GSR; Central: EEG; Audio, Visual, Depth |

**Table 2.** Non-emotional stimuli: Mean (sd) of valence and arousal ratings for non-emotional stimuli are presented. See whether it is needed or not

| Non-emotional | Valence | Arousal |
| --- | --- | --- |
| First | 4.59 (0.5) | 5.09 (0.69) |
| Second | 4.68 (0.83) | 4.9 (0.71) |

**Table 3.** Selected stimuli for EEG study from the stimuli dataset we have created (for stimuli^23^; for related article^13^). The time duration of each stimulus is 60 seconds.

| Stim Name | Target Emotion |
| --- | --- |
| ashayen (199) | Adventurous |
| horror (214) | Afraid |
| Anacondas_The_Hunt_for_the_Blood_Orchid_clip (10) | Alarmed |
| Lage_Raho_Munnabhai_Only_the_Funny_Scenes_[360p]_2 (99) | Amused |
| anger_legend_of_baghat_singh (198) | Angry |
| Divergent_Kiss_scene_clip (54) | Aroused |
| Butterfly_Nets (40) | Calm |
| Best_Horror_Kills_Ghost_Ship_Opening_scene (26) | Disgust |
| Jai_Ho (92) | Enthusiastic |
| SaddaHaq_1 (152) | Excited |
| MASOOM (109) | Happy |
| cheerfulRang_1 (201) | Joyous |
| Crash_Saddest_scene (51) | Melancholic |
| hate_lbs (210) | Miserable |
| Madari_movie_of_best_scene_[360p] (113) | Sad |
| Final_Race_of_Milkha_Singh_Career_[360p] (67) | Triumphant |

### 1.5 Scales

To rate the scales (shown in the figure-1) and provide emotional categories (during the second stage) following scale related instructions were provided.

1. **Valence:**The dimension of valence is the negative-positive valence of the emotion and ranges from unpleasant to pleasant (unhappy-happy). This would be a continuous scale ranging from 1 to 9 with anchors unpleasant (1) to neutral (5) to pleasant (9). You can perform a mouse click at any place on the scale, and adjust the ratings by dragging a triangle marker which will appear upon your first click on the scale. On the bottom of the scale, the score corresponding to the place of the triangle marker will replace the text key (to know your rated score on the scale). Once you are confirmed about your ratings you can click on this button to proceed to the next scale. However, you are provided an upper time limit of ten seconds to respond. Cartoon pictures with animated facial expressions from unpleasant to pleasant emotions are provided above the rating scale to help you rate your responses.
2. **Arousal:**Physiological arousal is the activation state of emotion, which ranges from inactive to active state. Arousal will also be a continuous scale ranging from 1 to 9 with anchors inactive (1) to active (9). With this scale you will interact similarity as you were doing with the valence scale. For arousal scales also cartoon pictures animating inactive to active state are provided above the rating scale to help you rate your responses. Once you are confirmed about your arousal ratings, click the button to go to the next scale.
3. **Dominance:**A person’s feeling of dominance in a situation is based on the extent to which he/she feels unrestricted or free to act in a variety of ways. For example, imagine yourself sitting in an exam hall and unprepared for the exam. You might feel submissive to this situation. On the contrary, if you are well prepared, you will feel dominant in dealing with the pressure of exams. Dominance scale is also a continuous scale varying from submissive (1) to dominant (9). To depict different degree of dominance a cartoon picture with varying sizes are provided. Small size to large size of the cartoon are animating the submissive to dominant feelings. You will interact with and record your ratings on dominance scale similarly as you did with the valence and arousal scales. Once you are confirmed about your ratings, click the button to go to the next rating scales. Or if the upper time limit is reached (5 seconds), you will be automatically navigated to the next scale.
4. **Liking:**How much do you like the video on a scale of 1 to 5? Please Don’t confuse liking with valence scales. You may like even the context eliciting the negative feelings. For example, if someone has a test of sad songs, he/she may like it. Liking simply asks if you like the stimulus or not which may vary depending upon your test for movies, songs and other factors. This is a discrete scale with values 1, 2, 3, 4 and 5. Least liking will be rated as 1, fair liking will be rated as 3 and much liking will be rated as 5. You can perform a mouse click near the places where you see a little white mark on the scale. You can see your rated score displayed on the button below. Upon clicking this button or after ten seconds, you will be navigated to the next scale.
5. **Familiarity:**Are you familiar with the content of the video? Rate on a scale of 1 (least) to 3 (fair) to 5 (much) based on your prediction of the next event in the stimulus you are looking at. For example, a scene of someone running and you are able to predict what could be the next event in the stimulus. Like the liking scale, familiarity is also rated on the discrete scale with discrete values 1, 2, 3, 4, and 5. You can click near the discrete points on the scale marked with a little white vertical bar. The rated score will be displayed on the button below the scale. Upon clicking this button or after five-second, you will be navigated to the next scale.
6. **Relevance:**Relevance scale varied from from not related (1) to completely related (5). It was a continuous scale. The relevance was assessed based on the assessment of any personal past life event that resembles the situation shown in the stimulus video.
7. **Emotion category:**The emotion categories are divided into four quadrants (as shown in the diagram). As per your ratings on the valence and arousal scale, you will be provided with a list of emotions in anyone quadrant. However, if you don’t find an emotion category in the list, you can enter the name of the emotion in the text box with the label “Propose New Emotion Category”.

**Figure 1.**
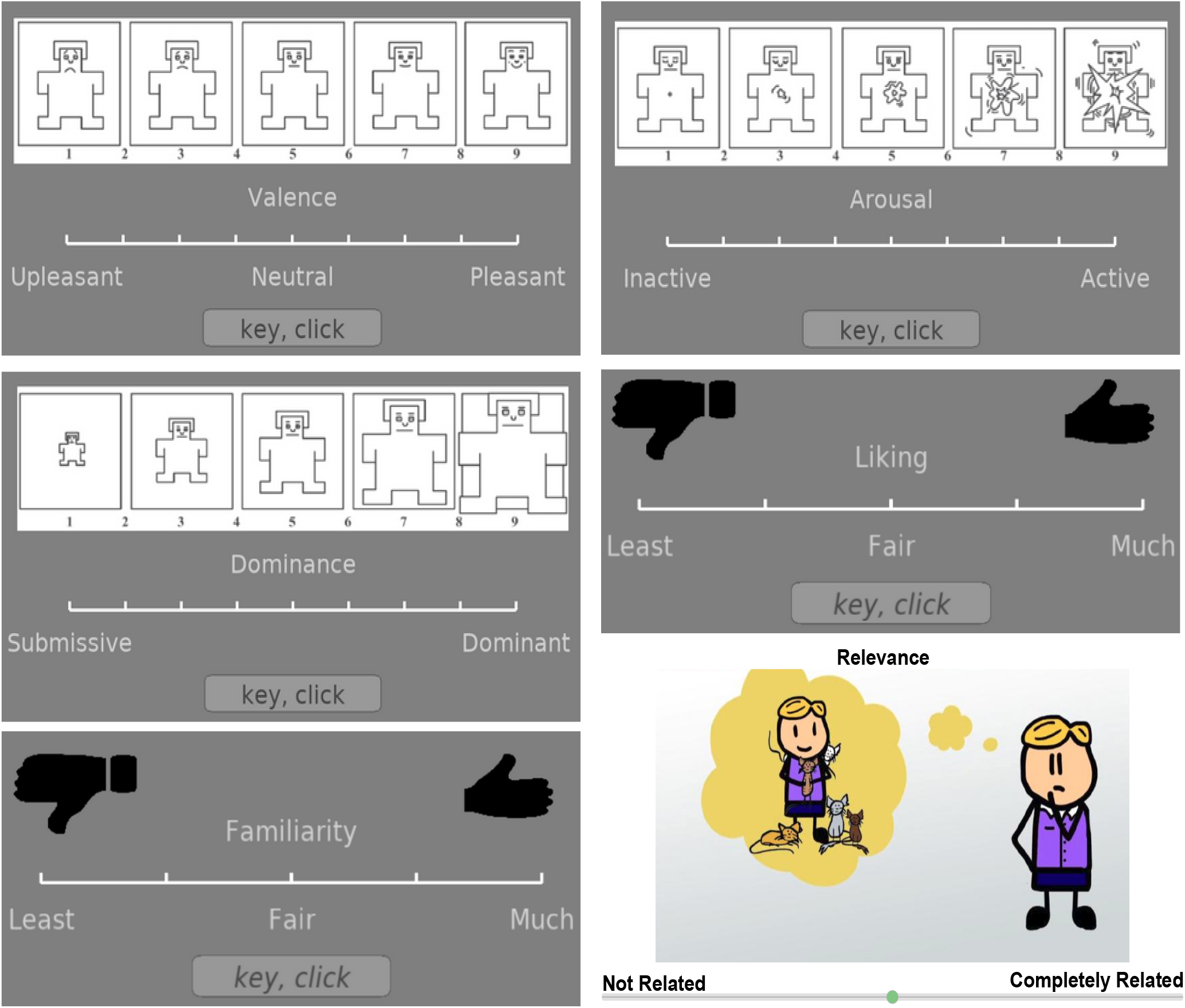
Scales used during the EEG experiment

### 1.6 Experiment Setup

#### 1.6.1 Laboratory Environment

The experiment was performed in a single laboratory environment. During the experiment, all the lights of the room were switched off. Participant set in front of a 15.6 computer screen (with resolution 800×600). Audio feedback were given using Sennheiser CX 180 Street II in-Ear Headphone. For response, a mouse and a keyboard were also placed on the table.

#### 1.6.2 EEG Setup

A 128 electrode system from EGI Inc. was used for the EEG recording. The raw data were stored in iMac using the NetStation recording software. In addition to EEG, peripheral signals including ECG and EMG were also recorded with Physio16 package. The sampling rate was 250 Hz.

### 1.7 EEG Experiment

#### 1.7.1 Experiment Overview

After providing consent, participants were provided instructions regarding the experiment and protocols using a presentation (experiment presentation [online]) by the experimenter. The EEG cap was placed correctly on the participant’s head. Electrodes were adjusted and refilled with the saline solution for conduction (if needed). The impedance was checked and kept under 30Kohm. In addition, peripheral channels ECG & EMG were correctly placed at the appropriate location on the body. The right and left ECG electrodes were placed directly below the clavicle near the right and left shoulder. The EMG electrodes were placed on the backside of the neck, left and right lateral gastrocnemius, and on the participant’s hand which was used for click. The participant performed a training/practice experiment for becoming familiar with the study. Questions from participants, if any were addressed after the practice session. Participants were instructed to call experimenter in case of any discomfort.

The experimenter continuously monitored the impedance after each trial (on the recording screen) while the participant responded on the response windows. If the impedance increased beyond the threshold, the experimenter paused the experiment, applied the saline solution to ensure that the impedance was below the threshold and resumed the recording again.

After completing the experiment, participants filled the google form and provided demographic details. We also asked for the time difference between when they were aware of felt emotion and when they clicked. Six participants did not respond. 30 out of 34 remaining participants reported that they clicked between 2-3 seconds after awareness for the feeling of emotion (fig-2a). In addition, we asked participants to rate their mood, cognitive load during the experiment (fig-2d), and how much they felt interfered by the clicking while watching the stimuli (fig-2c).

**Figure 2.**
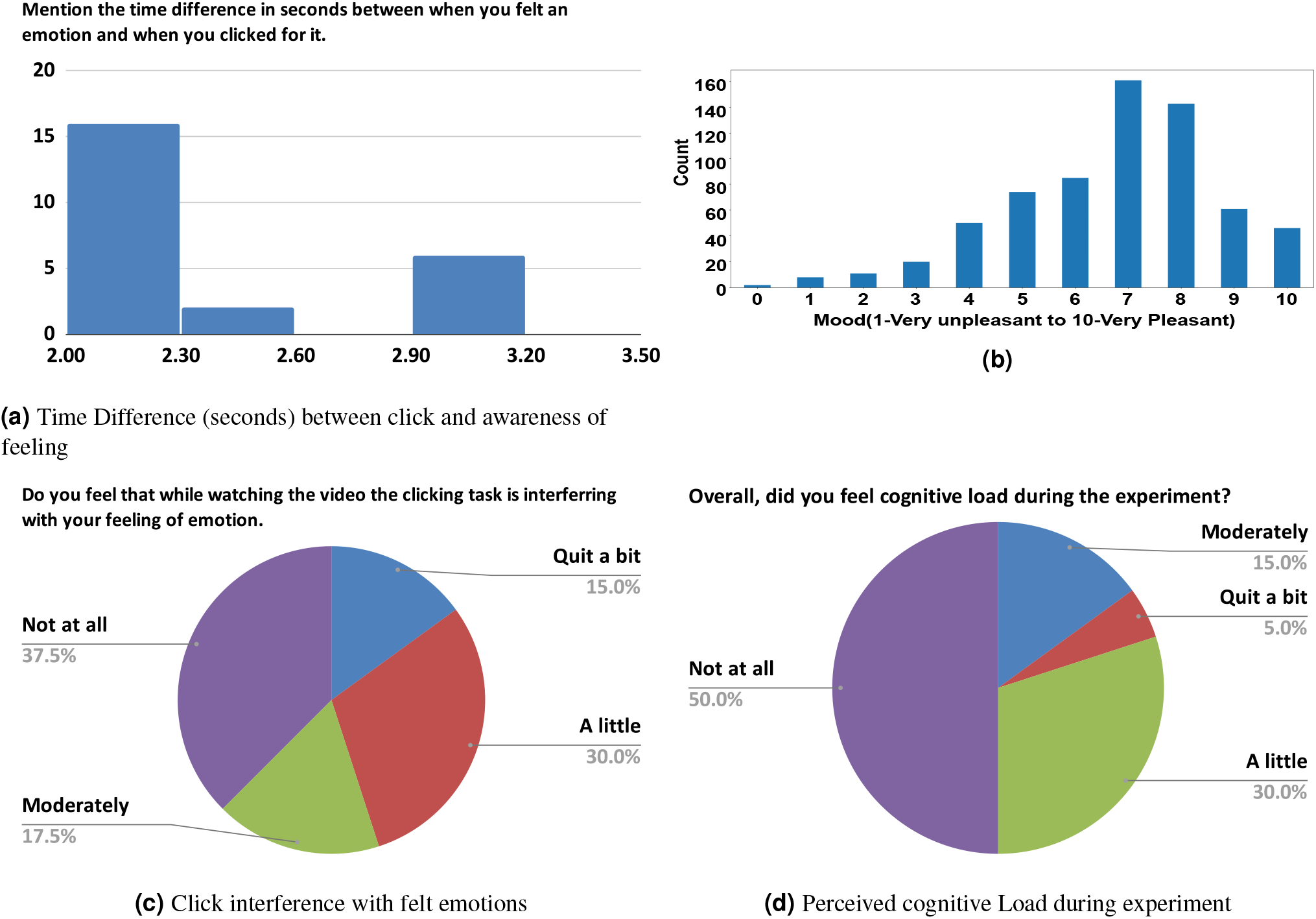
Participants Questionnaire. (a) Time difference between the moment when participants became aware about emotional feelings and the clicking, (b) overall general mood of the participants while performing the experiment, (c) the amount of interference of the non-emotional clicking task on felt emotions, and (d) the perceived cognitive load participants felt during the experiment.

#### 1.7.2 Experiment Paradigm

The paradigm comprised of bio-calibration, training, and testing modules.

##### Bio-calibration

The experiment in the laboratory starts with bio-calibration. Participants were asked to perform twenty specific tasks based on the instructions that appeared on the monitor screen. The tasks included Look right, Look left, Lookup, Look down, Wiggle your left toe, Hold your breath, Take a deep breath, Breath-in & breath-out, Grit teeth, Make the click event, Move hand slowly between keyboard and mouse, Press right arrow key in the keyboard, Imagine the legs movement on a beat, Imagine the finger movement on the beat, Look straight ahead, Move the eyeball left, right, up, down, Tongue movement, Chewing, Swallowing, Close the eyes. The physiological signals during these tasks were recorded to create a template and for possible use in reducing noise in the EEG signal (if considered part of the analysis).

Participants were trained first using three stimuli. During training the experimenter were present to assist in understanding the paradigm and response scales. After clearing all the doubts participants participated in the real experiment.

1. *Quiz:* We ran this module during training only. This sub-module assessed the participants’ understanding of the Valence and Arousal scales. A set of multiple-choice questions (stating an imaginary emotional scenario in English textual form on the screen; experiment presentation [online]) were displayed to the participants one by one (total 11 questions; experiment presentation [online]). After each question, participants had to select one option from the four given choices. The four choices were a) High Valence - High Arousal (HVHA), b) High Valence - Low Arousal (HVLA), c) Low Valence - High Arousal (LVHA), and d) Low Valence - Low Arousal (LVLA). The experiment proceeded to the next sub-module once the participants correctly answered at least 10 out of 11 questions.
2. *Resting-State Recording:* Participants were instructed to sit in a relaxing posture with no large physical movement. They were told not to click or use mouse for the duration of the appearance of the fixation mark on the screen (eyes opened). The duration of this baseline was 10 seconds during training and 80 seconds during real experiment.
3. *Stimuli Presentation & responses:* The practice/training module consisted of three trials. In each trial, participants were presented a 1-minute long stimulus video. Participants were instructed to click anywhere on the screen with the help of the mouse during the stimulus video presentation only when they felt any discernible emotion. Participants were instructed that they could click multiple times during the video. It is reported in the literature that the mouse clicking doesn’t interfere with the feelings of emotions^14^. The number of clicks were left to the choice of the participants with no lower or upper limit. The response window appeared once the stimulus video window closed.

In the set of response windows, after each stimulus video, participants were asked to rate valence, arousal, dominance, relevance, liking, and familiarity for the whole video. For valence, arousal, and dominance, participants had to choose a point on a continuous scale ranging from 1 to 9 with the help of the mouse. Manikins were also visible to assist the participants (as shown in the experiment presentation [online]). The valence, arousal, and dominance scales range from unpleasant to pleasant, inactive to active, and submissive to dominant, respectively. For liking and familiarity, participants had to choose a point on a discrete scale of range 1 to 5. Both the scales range from Least to Much. For Relevance, participants had to choose a point on a continuous scale ranging from 1 (Not related) to 5 (Completely related) spectrum.

After that, each click event had to be categorized in a particular emotion category selected from a list of presented emotions for each quadrant of valence-arousal space. For each click-event, a new window (click window; for reading convenience) appears after rating the scales mentioned above. In each click window, three consecutive images (the image frames at 20 frames before click, at click, 20 frames after click, from left to right) are shown on a single screen to aid the participants in recalling the context and the emotion they felt. In the bottom panel of the same window, the participants chose an emotion they felt while clicking. To choose an emotion, they first had to select a category among four categories-HVHA, HVLA, LVHA, LVLA and then selected an emotion from the presented list of emotions.

Participants could take as much time as they want to provide the ratings and classifications. Once the participants provided all the ratings and answered the questions for a stimulus video, the trial finished. A fixation mark was presented between trials. In between two stimulus videos, a simple distractor arithmetic task was given to the participants to shift their attention away from the emotional video seen earlier. Participants could take as much time as they wanted and they indicated their readiness for the next trials by a left mouse click. The subsequent trial began after the click.

During real experiment the procedure was exactly same with two exceptions. First, instead of three stimuli participants were shown 9 emotional stimuli. Second, during the real experiment participants were also shown non-emotional stimuli. One at the beginning and another between fifth and eight stimuli.

##### Overall subjective feedback

At the end of the experiment, subjects are asked to fill the google form asking for name, age, gender, and other details (Detail of Google Form [online]). Participants are also asked if they felt cognitive load during the experiment. The number of participants are responded in five categories as follows-not at all:20, a little:12, moderately:0, quit a bit:2, very much:0. Participants are asked, “Do you feel that while watching the video, the clicking task is interfering with your feeling of emotion. For example, you were more concerned about clicking than getting involved with the video and feel emotions.” The number of participants are responded in five categories as follows-not at all:15, a little:12, moderately:7, quit a bit:6, very much:0.

## 2 Quality Assurance (QA)

The physiological signals are recorded using EGI EEG 128 channel system (10/10 international systems for EEG electrode placement). In addition to EEG, ECG and EMG data was also recorded. We tried to maintain the impedance below 30 Kohm. The raw data for all the participants is assessed for head movement or whole body movement. Three participants based on this manual assessment were removed. The data are published in its original form to allow maximum flexibility in its use.

Artifacts from the Data was also removed as per the procedure described in associated publications. A very brief review of artifact removal we state here. Data is band pass filtered from 1-40 Hz followed by manual assessment and removal of detached electrodes (at any point out of whole recording duration) in the data. ICA is applied for the removal of other artifacts.

A unique quality of our data is that we provide the temporal localization of the emotional experience which had never been reported before in the emotion literature. In this dataset, participants were able to mark the time when they felt any emotion at the same time they are provided enough context for the emotional experience to evolve. Considering this aspect of our data, we claim that the dataset which we are providing is unique and can seed to new ideas in performing emotion experiments in lab setting without compromising the ecological validity.

## 3 Ethics

The complete experiment was conducted as per the guidelines of the Institutional Ethics Review Board, University of Allahabad (protocol code 2017-100 approved on Dec 8, 2017). All the content of the experiment was reviewed by the review board. Written informed consent was obtained from the participants.

## 4 Data Records

The dataset is available in a public repository openneuro^15^. The description of different files are as follows:

- Dataset_description: Describes the metadata for the dataset.
- participants.json and participants.tsv: Participants related details.
- self-assessment-feedback.tsv: Response by subjects on self-assessment scales, including valence, arousal, dominance, liking, familiarity, and relevance. Rating of emotional category is also provided in this file. The file also contains time-stamp of mouse click.
- Files with.fdt and.set extension provides raw EEG data.

## 5 Technical Validation

We performed technical validation on our dataset.

We validated the film clip dataset with 678 participants in two stages. We used sentiment analysis of YouTube comments and multimedia content analysis for the initial selection of candidate stimuli from the 1500 multimedia videos downloaded from YouTube against 355 affective words taken from affect word-net^16^. For the final set of stimuli the agreement among raters for self-assessment scales was calculated. The probability of elicitation of different emotional categories was also calculated and reported^13^.

The EEG and ECG data were validated by analyzing cardiac-brain activity during emotional experience^17^. In the cardiac-brain interaction analysis, linear and logistic mixed effect regression modeling was performed to assess how the degree of context-familiarity influences the cardiac-brain activity during the emotional experience.

Another analysis on EEG using dynamic functional connectivity and regression modeling was performed to assess in which frequency band the functional connectivity of different emotions was most distinct. And, how the reorganization of the functional connectivity is related with the different dimensions of emotional experience^18^. We found that emotional experiences differ in upper beta bands primarily and dynamics of functional connectivity is related with the arousal and dominance.

In addition we performed whole brain neural coordination analysis using microstate analysis method^19^. We observed different set of microstates for different emotional experiences. These microstates were source localized to brain regions responsible for socio-emotional processing and context comprehension. The sequence of transitions among the microstate also encode information related to different emotional experiences.

Using deep learning methods, we performed emotion recognition using the DENS dataset^20^. In one of the study we highlighted the importance of considering dominance dimension^20^. In another study we showed that capturing emotional events using our EEMP paradigm compared to other datasets^6,8,21^ increased the accuracy of emotional classification^22^.

